# Discovery of A Multi-Class Antibiotic Potentiator Against Resistant *Klebsiella pneumoniae*

**DOI:** 10.64898/2026.08.02.742369

**Authors:** S. L. Harris, S. Dutta, W. Thong, M. Morris, Z. Wang, X. Wang

**Affiliations:** Department of Chemistry, University of Colorado, Boulder, Colorado, 80309, USA

## Abstract

The rapid rise of multidrug-resistant (MDR) bacterial infections has severely limited treatment options, particularly for Gram-negative pathogens such as *Klebsiella pneumoniae*, a leading contributor to pneumonia, bloodstream, urinary tract, and surgical-site infections. One strategy to restore antibiotic efficacy is the use of resistance-mitigating agents (RMAs), compounds that re-sensitize bacteria to existing antibiotics without displaying independent antibacterial activity. Herein, we report the results of a high-throughput screen of a 3,200-compound fragment-based library against an MDR *K. pneumoniae* isolate in the presence of subinhibitory ciprofloxacin. This screen identified a tetrahydrocarbazole-containing compound, **1**, as a ciprofloxacin potentiator. Subsequent structure-activity relationship studies yielded a difluorinated analog, compound **5**, which potentiated multiple antibiotic classes in MDR *K. pneumoniae*, reducing MICs up to ≥16 fold. Further testing demonstrated synergistic interactions between compound **5** and ciprofloxacin, ceftriaxone, cefoxitin, and tetracycline across four genetically diverse MDR *K. pneumoniae* strains. These findings suggest that tetrahydrocarbazole-containing compounds constitute a promising new class of RMAs with potential for future development as therapies against MDR *K. pneumoniae* infections.

## INTRODUCTION

Antimicrobial resistance (AMR) is a growing global health threat. In 2021, bacterial AMR was associated with an estimated 4.71 million deaths worldwide, and current projections suggest it will be associated with 169 million deaths between 2025 and 2050 (1). Recent global surveillance data from the World Health Organization (WHO) further demonstrate the accelerating nature of the crisis, with one in six bacterial infections resistant to antibiotics in 2023, and resistance rising across 40% of pathogen-antibiotic combinations between 2018 and 2023 (2).

Gram-negative pathogens are particularly concerning due to the combined action of multiple intrinsic and acquired resistance mechanisms, including the outer membrane, multidrug efflux pumps, and β-lactamases (3). Among these organisms, *Klebsiella pneumoniae* is a leading cause of hospital-acquired bloodstream infections and a major contributor to pneumonia, urinary tract, and surgical-site infections (2, 4). It readily acquires and disseminates resistance determinants, driving the global spread of multidrug-resistant (MDR), extended-spectrum β-lactamase (ESBL)-producing, and carbapenem-resistant strains (5). According to the WHO, over 55% of *K. pneumoniae* bloodstream infections are resistant to third-generation cephalosporins, with increasing resistance to carbapenems and fluoroquinolones further narrowing treatment options (2).

Resistance-mitigating agents (RMAs, *a.k.a.*, antibiotic adjuvants) represent a promising strategy to combat AMR. These compounds re-sensitize bacteria to antibiotics without exhibiting significant independent antibacterial activity, thereby extending the lifespan of existing drugs and potentially reducing the selective pressure for resistance (6). This approach has been clinically validated by β-lactamase inhibitors, such as clavulanic acid (7), but there remains a need to identify RMAs that act through alternative mechanisms. Of particular interest are RMAs capable of potentiating multiple antibiotic classes. Such RMAs often act by increasing intracellular antibiotic accumulation, either through disruption of the outer membrane (8) or inhibition of multidrug efflux pumps (9), though few have advanced beyond preclinical development.

Bacterial two-component systems (TCSs) are an underexplored target for antibiotic potentiation. These signaling systems, composed of a sensor histidine kinase and a cognate response regulator, enable bacteria to sense and respond to environmental and antibiotic-induced stresses (10). The CpxRA TCS, which is highly conserved among Enterobacteriaceae including *Escherichia coli* and *K. pneumoniae*, was originally characterized as a response to misfolded proteins in the periplasm but is now recognized as a broader regulator of envelope homeostasis, influencing the expression of hundreds of genes involved in protein quality control, transport, cell wall synthesis, and other envelope-associated processes (11, 12). CpxA possesses both kinase and phosphatase activities that determine the phosphorylation state of the response regulator CpxR and, consequently, expression of the Cpx regulon.

Perturbation of Cpx signaling has been shown to alter susceptibility to several antibiotic classes, including β-lactams and aminoglycosides, although these effects are highly context dependent. For β-lactams, moderate activation of the Cpx response in *E. coli* promotes resistance, whereas overactivation causes growth, division, and morphological defects that increase susceptibility (13). In contrast, constitutive Cpx activation enhances survival during aminoglycoside exposure, although no single Cpx regulon member has been shown to account for this phenotype, suggesting that aminoglycoside resistance is influenced by the combined activity of multiple regulon members. (14). Thus, CpxRA represents a promising, but mechanistically incompletely understood, target for RMAs. Notably, although a putative CpxRA system has been identified in *Pseudomonas aeruginosa*, its regulon differs substantially from that of *E. coli* (15). Furthermore, no homologous system has been identified in *Acinetobacter baumannii* (16), indicating that CpxRA-directed potentiators may display species selectivity. Species-selective RMAs may nonetheless be highly valuable, as they enable targeted restoration of antibiotic activity in priority pathogens while minimizing effects on non-target bacterial populations.

Here, we report the discovery of a novel class of RMAs identified through a high-throughput screen of a 3,200-member fragment-based library against an MDR *K. pneumoniae* isolate. This screen identified compound **1** (Figure 1) as a ciprofloxacin potentiator. Compound **1** and related tetrahydrocarbazoles were previously reported to exhibit antibacterial activity in *E. coli* through proposed inhibition of the CpxA phosphatase, although their activity as RMAs has not been explored (17-19). Subsequent structure-activity relationship studies led to the identification of a difluorinated analog that exhibited significantly improved activity and potentiated multiple classes of antibiotics in *K. pneumoniae.* These findings establish tetrahydrocarbazole-containing compounds as promising antibiotic adjuvants and motivate further investigation into their mechanism of action and the impact of CpxRA signaling on antibiotic susceptibility.

**Figure 1.**
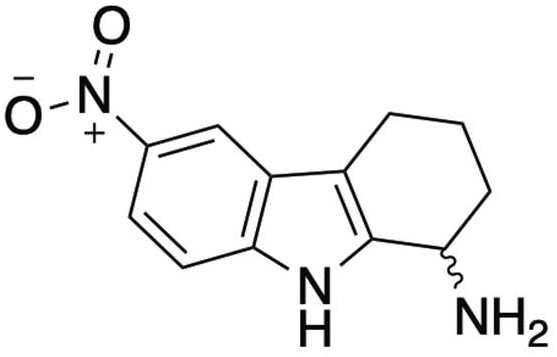
Structure of Compound **1**.

## RESULTS

The minimum inhibitory concentration (MIC) of ciprofloxacin (CIP) against the MDR *K. pneumoniae* AR-1022 strain was determined to be 0.5 µg/mL using the Clinical and Laboratory Standards Institute (CLSI) broth microdilution method (20). The AR-1022 strain was obtained from the CDC and FDA Antimicrobial Resistance Isolate Bank (21). CIP was chosen because of its widespread clinical use in the treatment of Gram-negative infections and the growing prevalence of fluoroquinolone resistance (2). Encouraged by our prior success discovering [1,2,5]oxadiazolo[3,4-*b*]pyrazine-containing colistin adjuvants from the TimTec fragment-based library, we conducted a high-throughput screen of this library in AR-1022 in the presence of subinhibitory CIP (0.25 µg/mL) (22, 23). The 3,200 compounds in this library have low molecular weights (110–290 Da) and relatively high aqueous solubility, making them well suited for screening against Gram-negative bacteria.

Compounds were screened at a final concentration of 7.5 µM in 96-well plates containing cation-adjusted Mueller-Hinton broth supplemented with 0.25 µg/mL CIP, corresponding to the clinical breakpoint in Enterobacteriaceae. The screen was multiplexed, with compounds from two library plates combined into a single screening plate, after which hits were deconvoluted and verified individually. Following incubation at 37°C for 18 hours, eight compounds inhibited visible bacterial growth in combination with CIP (Figure S1). These hits were selected for confirmation using the same assay format, with each compound tested at 30 µM, 15 µM, and 7.5 µM in the presence and absence of 0.25 µg/mL CIP. This approach identified the compound 6-nitro-2,3,4,9-tetrahydro-1H-carbazol-1-amine (**1**, Figure 1) as a potentiator of CIP. Compound **1** was prioritized for further study over the other seven hit compounds due to its RMA activity rather than direct antibacterial effects, as well as its structural divergence from CIP.

Checkerboard analysis in AR-1022 demonstrated that **1** was synergistic with CIP, with a fractional inhibitory concentration index (FICI) of 0.375, below the established threshold for synergy (≤0.5). **1** has previously been reported as an antibacterial agent in *E. coli* by van Rensburg et al., Li et al., and Fortney et al., however, it has not been characterized as an antibiotic adjuvant (17-19). Prior studies have further suggested that **1** and related analogs function as inhibitors of the CpxA phosphatase. Given that the CpxRA system is conserved across Enterobacteriaceae, we hypothesized that inhibition of this pathway may contribute to the observed RMA activity in *K. pneumoniae*. To investigate this, four representative analogs of **1** described by Li et al. were synthesized (Table 1) to assess whether structure-activity relationships reported in *E. coli* would be recapitulated in *K. pneumoniae* and to identify derivatives with enhanced potency (18). **1** and its analogs were synthesized according to established procedures described by Li et al.

**Table 1.** Structure-activity relationship studies of compound **1** analogs.

| Compound | R<sub>1</sub> | R<sub>2</sub> | MIC<br>(μg/mL) | CIP MIC<br>(μg/mL)<sup>1</sup> | Fold of<br>Potentiation<sup>2</sup> | GI<sub>50</sub><br>(μg/mL) |
| --- | --- | --- | --- | --- | --- | --- |
| **1** | NO<sub>2</sub> | H | 32 | 0.25 | 2 | 12 |
| **2** | H | H | >64 | 0.5 | 1 | 68 |
| **3** | Cl | H | 2 | n.d. | n.d. | 7.0 |
| **4** | F | H | 64 | 0.125 | 4 | 9.7 |
| **5** | F | F | 32 | 0.0625 | 8 | 8.5 |

<sup>1</sup>In the presence of 4 μg/mL compound indicated.
<sup>2</sup>Calculated as the ratio of the ciprofloxacin MICs in the absence and presence of 4 μg/mL compound.
n.d.: Not determined.

The MICs of the analogs alone were determined in AR-1022 as described previously. **1**, **2**, **4**, and **5** exhibited limited intrinsic antibacterial activity, with MIC values ranging from 32 to >64 µg/mL (Table 1). In contrast, the chloro-substituted analog, **3**, displayed increased antibacterial activity, with an MIC of 2 µg/mL. To assess potentiation, the MIC of CIP was tested in combination with 4 µg/mL of each compound in AR-1022, excluding **3** due to its low MIC value. Of the analogs tested, the fluorinated derivatives showed the greatest enhancement of CIP activity. The difluorinated derivative, **5**, reduced the CIP MIC 8-fold to 0.0625 µg/mL, while the monofluorinated derivative, **4**, reduced the MIC 4-fold to 0.125 µg/mL. In contrast, the unsubstituted analog, **2**, demonstrated no RMA activity. The mammalian toxicity of these compounds was then evaluated in human hepatocellular carcinoma (HepG2) cells using CellTiter-Glo viability assays (Promega). The half-maximal inhibitory concentrations (GI_50_s) were calculated using GraphPad Prism. **1** and its analogs generally exhibited GI_50_ values in the range of 7-12 µg/mL, with **5** showing a GI_50_ of 8.5 µg/mL. This value is more than two-fold higher than the concentration required for CIP potentiation in AR-1022.

Given its strong potentiation of CIP and low independent antibacterial activity, compound **5** was selected for expanded antibiotic combination testing in *K. pneumoniae* AR-1022. The MICs of 13 antibiotics representing 7 distinct classes (fluoroquinolones, cephalosporins, carbapenems, aminoglycosides, tetracyclines, rifamycins, and polymyxins) were tested alone and in combination with 4 µg/mL **5** in AR-1022 (Table 2). Compound **5** potentiated multiple antibiotic classes, resulting in at least a 2-fold reduction in MIC for every antibiotic tested. The greatest effect was observed with ceftriaxone, for which the MIC decreased from >512 µg/mL to 32 µg/mL in the presence of **5**, corresponding to a ≥16-fold reduction. Susceptibility to CIP improved 8-fold with the addition of **5**, while ofloxacin, levofloxacin, cefoxitin, tetracycline, and rifampicin each demonstrated 4-fold MIC reductions. 2-fold reductions were observed for imipenem, tobramycin, and colistin. These results suggest that **5** acts as a multi-class antibiotic potentiator capable of sensitizing *K. pneumoniae* to mechanistically distinct antibacterial agents.

**Table 2.**
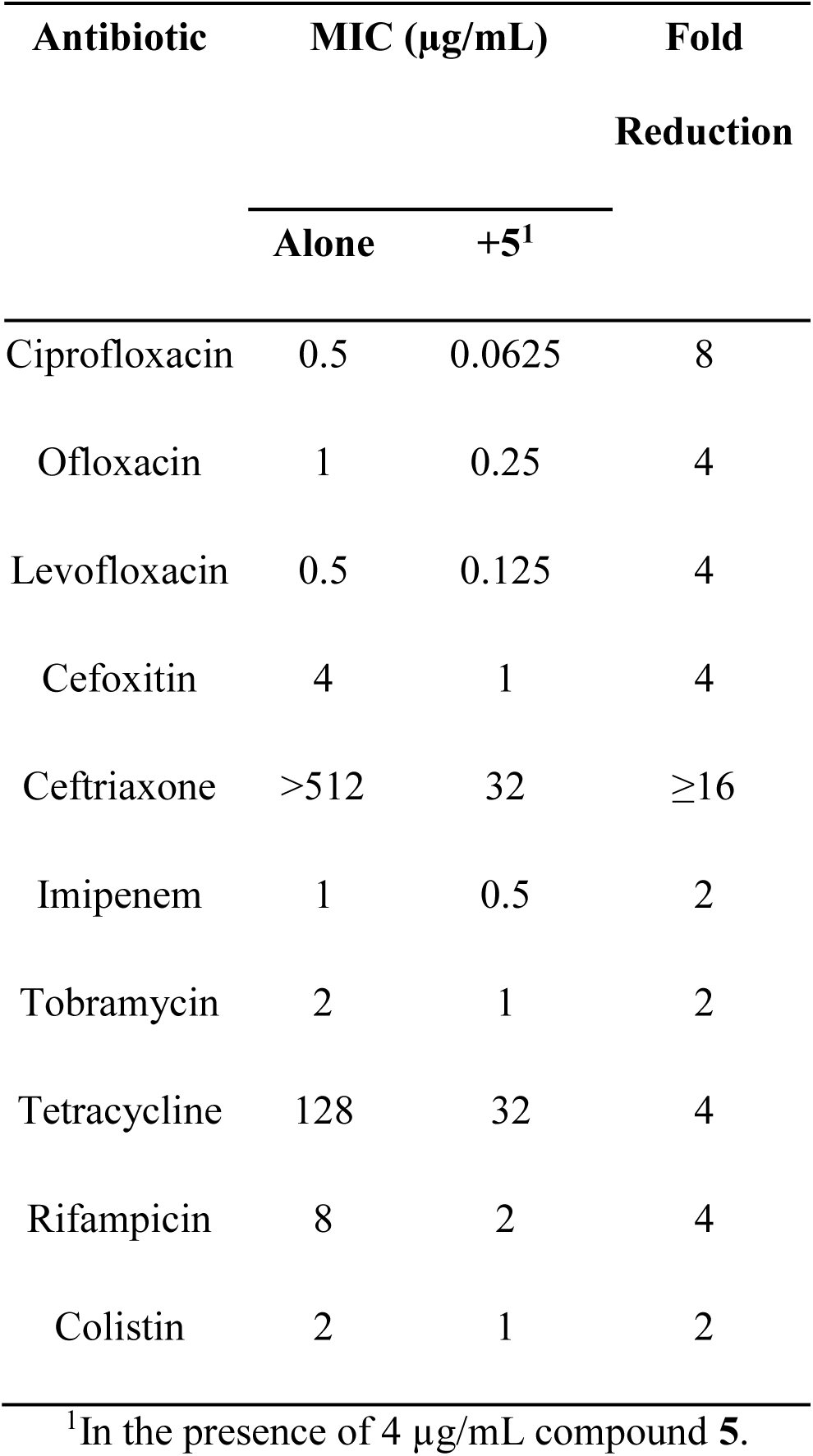
Evaluation of **5** with various antibiotics in *K. pneumoniae* AR-1022.

| Antibiotic | MIC (µg/mL) |  | Fold<br>Reduction |
| --- | --- | --- | --- |
|  | Alone | +**5**<sup>1</sup> |  |
| Ciprofloxacin | 0.5 | 0.0625 | 8 |
| Ofloxacin | 1 | 0.25 | 4 |
| Levofloxacin | 0.5 | 0.125 | 4 |
| Cefoxitin | 4 | 1 | 4 |
| Ceftriaxone | >512 | 32 | ≥16 |
| Imipenem | 1 | 0.5 | 2 |
| Tobramycin | 2 | 1 | 2 |
| Tetracycline | 128 | 32 | 4 |
| Rifampicin | 8 | 2 | 4 |
| Colistin | 2 | 1 | 2 |

<sup>1</sup>In the presence of 4 µg/mL compound **5**.

The activity of **5** was evaluated in additional MDR *K. pneumoniae* strains in combination with CIP, ceftriaxone (CRO), cefoxitin (FOX), and tetracycline (TET) using checkerboard broth microdilution assays (Table 3). CIP and CRO were selected for having the largest MIC fold reductions in the preceding dataset, while FOX was included to further probe β-lactam potentiation. Although several antibiotics exhibited 4-fold MIC reductions, TET was chosen because it represents a mechanistically distinct class from fluoroquinolones and β-lactams and is more commonly used in Gram-negative bacteria than rifampicin. Across all strains tested, **5** had a consistent MIC of 32 µg/mL. Synergy with CIP also remained consistent, with FICI values of 0.25 observed in *K. pneumoniae* AR-1013, AR-0497, AR-1022, and AR-1028, all obtained from the CDC and FDA Antimicrobial Resistance Isolate Bank (21). Synergy was also observed with CRO across all four strains, with FICI values ranging from ≤0.14 to 0.38. FOX combinations similarly exhibited synergistic interactions, with FICI values between 0.19 and 0.50. TET combinations yielded FICI values of 0.38–0.50 in *K. pneumoniae* strains AR-1013, AR-1022, and AR-1028; however, the FICI for AR-0497 could not be determined due to the limited aqueous solubility of TET, which led to precipitation at higher concentrations and prevented reliable interpretation. Collectively, these findings indicate that **5** retains RMA activity across genetically distinct *K. pneumoniae* isolates and can potentiate multiple classes of antibiotics.

**Table 3.**
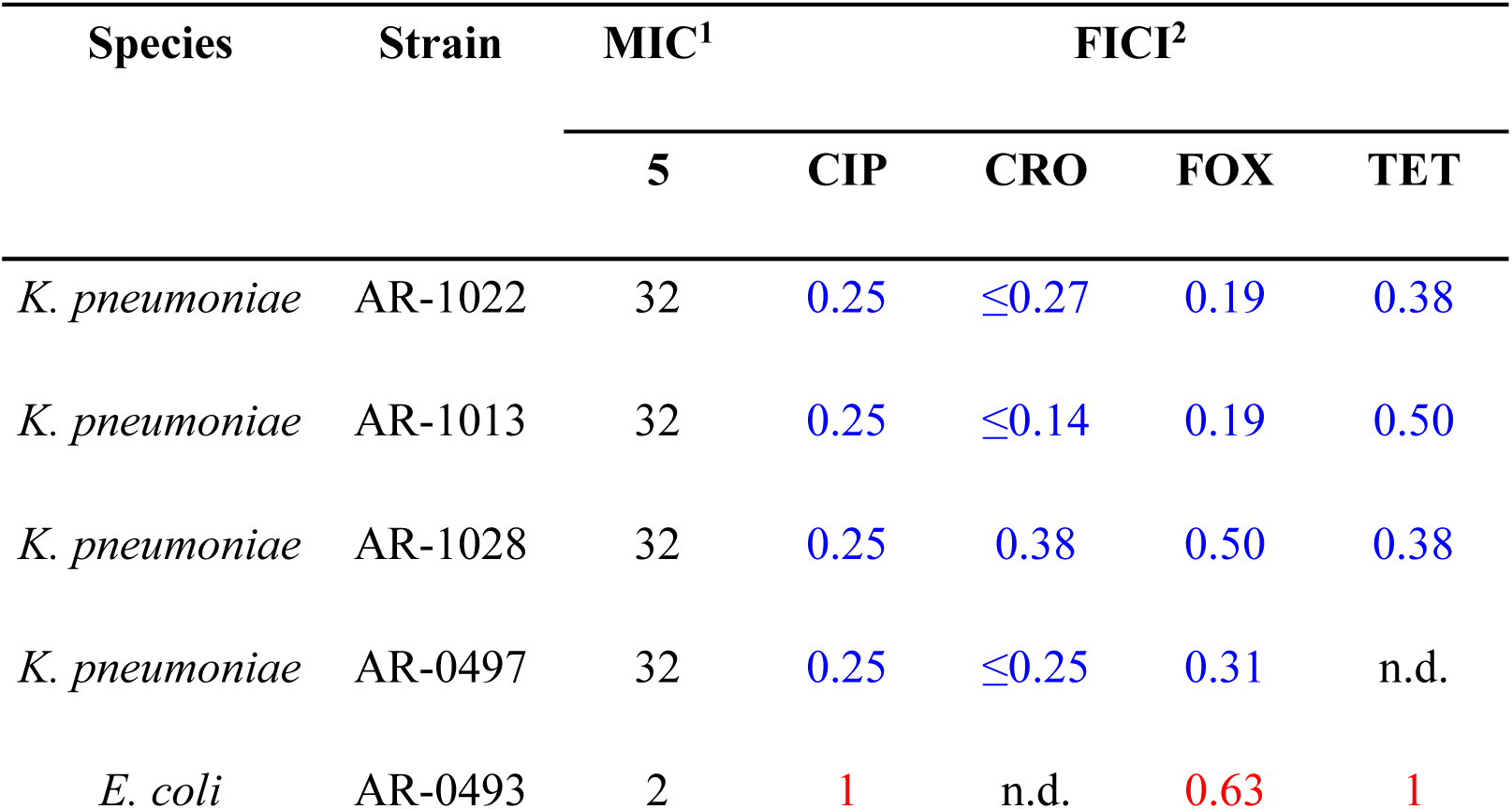
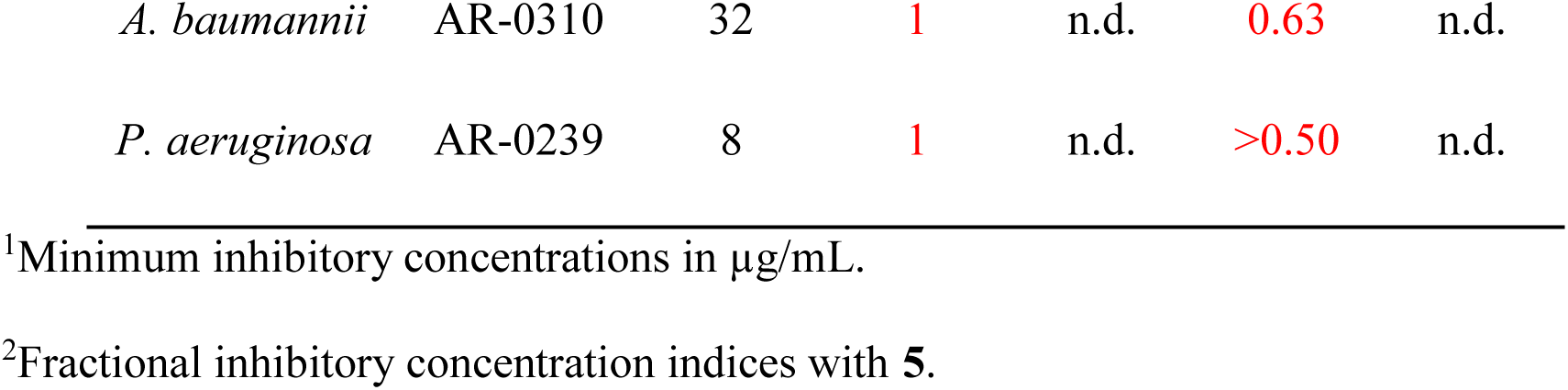
Results of checkerboard broth microdilution assays with **5** and CIP, CRO, FOX, or TET.

The activity of **5** was next characterized in additional Gram-negative bacterial strains acquired from the CDC and FDA Antimicrobial Resistance Isolate Bank (21). In *E. coli* AR-0493, **5** had a low MIC of 2 µg/mL; however, combination testing with CIP, FOX, and TET yielded no evidence of synergy, with FICI values of 1, 0.63, and 1, respectively (Table 3). Similarly, in *A. baumannii* AR-0310 and *P. aeruginosa* AR-0239, combinations with CIP, FOX, and TET failed to demonstrate synergy, with FICI values >0.5 observed for both organisms where testing was feasible. Together, these results suggest that the RMA activity of **5** is restricted to *K. pneumoniae*, although independent antibacterial activity exists in *E. coli*.

## DISCUSSION

Compound **5** enhanced the activity of mechanistically distinct antibiotics, including fluoroquinolones, cephalosporins, tetracycline, and rifampicin, and retained synergistic activity across all MDR *K. pneumoniae* strains tested. This broad potentiation profile suggests that **5** affects a general bacterial process influencing antibiotic susceptibility rather than directly targeting a single antibiotic-specific mechanism.

Previous studies proposed that compounds within this class inhibit the CpxA phosphatase, resulting in dysregulation of the CpxRA envelope stress response system in *E. coli* (17-19). CpxRA regulates multiple envelope-associated processes that can influence antibiotic susceptibility, making it a plausible mediator of compound **5**’s observed RMA activity in *K. pneumoniae* (11, 12). The agreement between the structure-activity relationships in this study and those previously reported in *E. coli* further supports this hypothesis. In both organisms, fluorinated derivatives exhibited the strongest activity, with the difluorinated analog, **5**, demonstrating the greatest activity.

Although compound **5**’s RMA activity was restricted to *K. pneumoniae*, this selectivity may be therapeutically advantageous. Species-targeted potentiators could enable more precise re-sensitization of high-risk pathogens while minimizing disruption to other bacterial populations. Similar to narrow-spectrum antibiotics, these RMAs may reduce selection for resistance and better preserve the host microbiome, which can experience long-term perturbations following broad-spectrum antibiotic exposure (24). The species selectivity observed in this study is also consistent with the CpxRA hypothesis. Although CpxA and CpxR are highly conserved between *E. coli* and *K. pneumoniae* (CpxA: 94.7% identity, 98% similarity; CpxR: 94.8% identity, 97.4% similarity) (25, 26), the *P. aeruginosa* Cpx system includes the *Pseudomonas*-specific adaptor proteins CpxM and CpxH and exhibits regulatory differences from the Enterobacteriaceae CpxRA system (15). Notably, *A. baumannii* lacks identifiable CpxA/CpxR homologs and instead relies on alternative systems, such as BfmRS and BaeSR, to mediate envelope stress responses (16). This may explain why activity was observed in *E. coli* and *K. pneumoniae* but not in the other two species. Interestingly, compound **5** demonstrated intrinsic antibacterial activity in *E. coli* AR-0493 but did not exhibit synergistic interactions with antibiotics in the same strain.

Despite the plausibility of CpxRA involvement, alternative mechanisms of action cannot be excluded. Many multi-class antibiotic adjuvants function through membrane permeabilization or efflux pump inhibition, both of which could account for the broad sensitization observed (8, 9). Additional studies to validate CpxRA involvement and assess effects on membrane permeability and efflux activity will therefore be necessary to establish the mechanism of action of this class of compounds.

Cytotoxicity studies in HepG2 cells demonstrated moderate mammalian toxicity for compound **1** and its analogs, with GI_50_ values generally ranging from 7-12 µg/mL. Although these values imply a somewhat narrow therapeutic window, compound **5** retained activity at concentrations greater than 2-fold below its GI_50_ value. These findings suggest that further optimization may improve selectivity while preserving RMA activity. Compound **5** was also evaluated by Fortney et al. in a murine model of uropathogenic *E. coli* infection, where it exhibited low *in vivo* toxicity (19). Treatment significantly reduced bacterial recovery from urine and showed trends toward reduced bacterial burdens in the bladder and kidneys. Collectively, these results indicate that compounds within this class have a favorable preliminary safety profile and sufficient therapeutic potential to warrant further optimization and mechanistic investigation.

## MATERIALS AND METHODS

### Compound Synthesis

All synthetic compounds were synthesized according to established procedures (18). Compound identity and purity were confirmed by ^1^H NMR spectroscopy and HPLC analysis, with all compounds demonstrating >95% purity.

### Bacterial Strains and Growth Media

All bacterial strains were obtained from the CDC and FDA Antimicrobial Resistance Isolate Bank (21). Strains were streaked onto tryptic soy agar plates supplemented with 5% sheep blood and incubated at 35°C for 18-24 hours.

### Antimicrobial Susceptibility Testing

Minimum inhibitory concentration experiments were performed according to the Clinical and Laboratory Standards Institute (CLSI) broth microdilution method (20). Bacteria were cultured in Luria-Bertani (LB) medium to the log phase and then diluted to an OD_600_ of 0.002 in cation-adjusted Mueller Hinton Broth (MHBII). Two-fold serial dilutions were prepared in clear 96-well plates (Life Science Products) by dispensing 100 µL of MHBII into row A and 50 µL into rows B-H. Antibiotics or compounds were added to row A at the desired starting concentrations, followed by two-fold serial dilutions down the plate using a 200 µL multichannel pipette. Wells were then inoculated with 50 µL of the bacterial suspension to achieve a final OD_600_ of 0.001. Plates were incubated at 35°C with shaking for 18 hours. The MIC was defined as the lowest concentration of compound that inhibited visible bacterial growth.

### Checkerboard Assays and FICI Determination

Checkerboard assays were performed according to the CLSI broth microdilution method (20). Bacteria were cultured in Luria-Bertani (LB) medium to the log phase and then diluted to an OD_600_ of 0.002 in cation-adjusted Mueller Hinton Broth (MHBII). Two-fold serial dilutions were prepared in clear 96-well plates (Life Science Products). Compound A (the antibiotic) was serially diluted two-fold along the horizontal axis (columns), and Compound B (compound **5**) was serially diluted two-fold along the vertical axis (rows). Wells were then inoculated with 50 µL of the bacterial suspension to achieve a final OD_600_ of 0.001. Plates were incubated at 35°C with shaking for 18 hours. Fractional inhibitory concentration (FIC) indices were calculated using the following formula: FIC of Compound A = MIC of A in combination / MIC of A alone; FIC of Compound B = MIC of B in combination / MIC of B alone; FICI = FIC(A) + FIC(B).

### Mammalian Cytotoxicity Assays in HepG2 Cells

Human hepatocellular carcinoma HepG2 cells (ATCC) were seeded in white, tissue-culture-treated 96-well plates (Corning 3917) in Dulbecco’s modified Eagle’s medium (DMEM) supplemented with 10% fetal bovine serum (FBS) and 1% penicillin/streptomycin at a density of 20,000 cells per well. The final medium volume in each well was 100 µL. Cells were incubated at 37°C in 5% CO_2_/95% air for 16 hours. The medium was then removed and replaced with 99 µL of fresh prewarmed medium. Subsequently, 1 µL of each compound in DMSO at the appropriate concentration was added to each well. After 24 hours of incubation at 37°C, plates were equilibrated to room temperature for 30 minutes. Then, 100 µL of CellTiter-Glo reagent (Promega) was added to each well and mixed for 2 minutes on an orbital shaker. Plates were incubated at room temperature for an additional 10 minutes, after which luminescence was measured using an EnVision Multilabel Plate Reader (PerkinElmer). GI_50_ values were calculated as half-maximal inhibitory concentrations using Prism (GraphPad).

## ACKNOWLEDGEMENTS

The authors thank the Venture Partners at the University of Colorado Boulder and the Colorado Office of Economic Development and International Trade (OEDIT) Advanced Industries Program for financial support.

